# Trans2express – de novo transcriptome assembly pipeline optimized for gene expression analysis

**DOI:** 10.1101/2024.01.11.575187

**Authors:** Aleksandra M. Kasianova, Aleksey A. Penin, Mikhail I. Schelkunov, Artem S. Kasianov, Maria D. Logacheva, Anna V. Klepikova

## Abstract

**Background:** As genomes of many eukaryotic species, especially plants, are large and complex, their *de novo* sequencing and assembly is still a difficult task despite progress in sequencing technologies. An alternative to genome assembly is the assembly of transcriptome, the set of RNA products of the expressed genes. While a bunch of *de novo* transcriptome assemblers exists, the challenges of transcriptomes (the existence of isoforms, the uneven expression levels across genes) complicates the generation of high-quality assemblies suitable for downstream analyses.

**Results:** We developed Trans2express – a web-based tool and a pipeline of *de novo* hybrid transcriptome assembly and postprocessing based on rnaSPAdes with a set of subsequent filtrations. The pipeline was tested on *Arabidopsis thaliana* cDNA sequencing data obtained using Illumina and Oxford Nanopore Technologies platforms. The comparison of structural characteristics of the transcriptome assembly with reference *Arabidopsis* genome revealed the high quality of assembled transcriptome with 86.1% of *Arabidopsis* expressed genes assembled as a single contig. We tested the applicability of the transcriptome assembly for gene expression analysis and showed high congruence of gene expression levels and sets of differentially expressed genes between analyses based on genome and based on the transcriptome assembly.

**Conclusions:** We present Trans2express – a protocol for *de novo* hybrid transcriptome assembly aimed at recovering of a single transcript per gene. We expect this protocol to promote the characterization of transcriptomes and gene expression analysis in non-model plants and web-based tool to be of use to a wide range of plant biologists.

## Background

Despite great progress in sequencing technology and decrease of sequencing costs, large-scale characterization of complete genomes still remains challenging and resource consuming. However, for many types of biological studies complete genome assembly is not necessary, and the data on the sequence of expressed genes RNA products (transcriptome) is sufficient. These directions of research include (but not limited to) the search of genes involved in traits of interest and their functional validation [1,2], the investigation of the evolution of gene families and genetic networks [3,4], the search of horizontal gene transfer [5], phylogenetic and population genetic studies [6,7]. The availability of transcriptome sequences also facilitates genetic studies enabling the development of markers for genetic mapping [8,9] and design of guide RNA for genetic editing.

*De novo* transcriptome assembly is usually carried out by shotgun sequencing with short read technologies [10–12]. In this case the typical primary outcome of *de novo* assembly is the high number (several hundred thousand) of relatively short contigs (N50 ∼ 1 kb) most of which represent only partial sequence of a transcript [13,14]. At the same time, most applications outlined above require one sequence per gene (ideally, full-length transcript of the isoform with the longest CDS). In order to reduce the number of contigs several stages of filtration and/or clustering are usually done [10,15]. This approach is error-prone and may result in missing transcripts by filtering them based on length or by collapsing (which is especially critical for polyploids). Many applications, especially ones dealing with quantitative analysis, are sensitive to these errors. In particular, it was shown that the use of short read-based assemblies for differential expression analysis leads to both false negative (the non-detection of differential expression, DE) and false positive (false detection of DE) errors [16].

Long reads generated by third-generation technologies such as PacBio and Oxford Nanopore Technologies (ONT) have potential to overcome these limitations because they allow to get complete transcript sequences. However, the accuracy of these technologies is still low (up to 10% of sequencing errors [17]). Moreover, the predominant type of errors are insertions and deletions which can affect ORF prediction [18]. Thus, by now the most prospective approach for the *de novo* assembly is the combination of short but accurate Illumina reads and long reads from PacBio or ONT [19]. It is widely used for genomes; there exists a number of tools for hybrid genome assembly (e.g. SPAdes [20], MaSuRCA [21], Unicycler [22]).

Transcriptome assembly however has its own challenges (the existence of isoforms, the uneven expression levels across genes), which makes them unsuitable and calls for the development of new protocols. By now the number of hybrid *de novo* transcriptome assemblers is limited and includes IDP-denovo [23] and rnaSPAdes [24]. IDP-denovo algorithm is a multistep procedure that includes 1) assembly using short reads alone; 2) error correction of long reads using mapping of short reads; 3) mapping of corrected long reads on short reads-alone assembly and scaffolding of short reads contigs. The obvious limitation of this approach is the accuracy of long reads. The authors of the algorithm tested it on PacBio reads which are more accurate than ONT and have a more random error profile [17]. Though error correction using information from short reads greatly improves accuracy of long reads generated by PacBio [25], for ONT long reads correction is not that efficient due to multiple indels [18]. Besides this, recent benchmarking of error correction algorithms shows that it generates a number of artifacts, in particular removal of minor isoforms and lowly expressed genes and distortion of multigene families [26]. Also, IDP-denovo is highly demanding for computational resources.

An alternative approach is the use of long reads for the resolution of a graph obtained from the assembly of short reads. The program that realizes this approach is rnaSPAdes [24,27]. SPAdes is a family of assembly tools which includes modules for standard genome, metagenome, transcriptome and single cell genome data. Specialized module for transcriptome assembly, rnaSpades, is an incorporation of a hybrid genome assembler hybridSPAdes [28] into a transcriptome assembly pipeline. rnaSPAdes allows the identification of potential gene isoforms and combination them into groups, reducing the transcriptome ambiguity, however, the number of resulting transcripts is still much higher than the number of genes [24].

We suggest a pipeline of *de novo* assembly of a transcriptome optimized for downstream gene-centered analyses – Trans2express. The pipeline includes *de novo* assembly using rnaSPAdes and several subsequent filtrations aiming to reduce the number of contigs per gene (these include incompletely assembled transcripts and isoforms). In order to test the efficiency of this approach, we sequenced and assembled the transcriptome of *Arabidopsis thaliana*, a model object of plant genetics with a very well characterized genome. We assembled and functionally annotated the transcriptome without reliance on the reference genome, and then compared the results with ones based on the genome. Since the goal of many transcriptome projects is the estimation of gene expression levels, we also compared the results of the analysis of differential gene expression based on reference and on *de novo* transcriptome assembly.

Trans2express is also available as an online tool (http://83.149.250.217/trans2express/) with easy-to-use interface.

## Methods

### Plant growth

*A. thaliana* (Columbia accession, CS70000) was grown in a climate chamber (16 hours light/ 8 hours dark, 22□, 60% relative humidity) for 21 days after germination. For transcriptome assembly mature leaves were collected in three replicates (each replicate represented a single leaf from a single plant) at ZT10 (Zeitgeber 10, ten hours after turning the light on). For differential expression analysis mature leaves were harvested at ZT09 (light) and ZT21 (dark) in two replicates; each replicate consists of a single leaf from seven plants.

### Sample preparation and sequencing

RNA was extracted using Qiagen RNeasy Mini kit (Venlo, Netherlands). RNA integrity was assessed using capillary electrophoresis on Agilent Bioanalyzer2100 (Santa Clara, CA, USA). NEBNext Ultra II RNA directional kit was used for Illumina library preparation (New England Biolabs, Ipswich, MA, USA) following manufacturer instructions. Libraries were sequenced on HiSeq4000 (Illumina, San Diego, CA, USA) in a paired mode with read length 150. For ONT libraries RNA was converted to cDNA using Mint cDNA synthesis kit (Evrogen, Moscow, Russia) with 18 cycles of amplification with the following modification: cDNA synthesis is primed with oligonucleotides that contain not only dT part and adapter part but also custom barcode, specific for each sample. Amplified cDNA was used as input for library preparation using standard protocol for genomic DNA with LSK-109 kit. Sequencing was performed on MinION with 9.4.1 flow cell.

### Illumina read processing

Raw Illumina reads were trimmed using fastp 0.23.4 [29] with following parameters: “-l 50 --cut_front --cut_right “.

### ONT read processing

Basecalling was performed by Guppy 3.4.5 (Oxford Nanopore Technologies, Oxford, United Kingdom); reads with mean quality less than 7 were removed (“*--qscore_filtering --min_qscore 7*”). After that, reads were processed by a custom pipeline NTproc (https://github.com/shelkmike/NTproc). NTproc performs the following operations consecutively:

1. NTproc trims PCR adapters.
2. NTproc removes all reads that do not have PCR adapters on both ends. This allows to eliminate reads coming from cDNAs that were fragmented after the reverse transcription or were not sequenced completely.
3. NTproc trims remaining (non-PCR) adapters.
4. NTproc makes reverse-complements of reads that have barcodes on 5’ ends. Since barcoded primers bind to 3’ ends of RNAs, this operation makes all reads oriented in the 5’→3’ direction of their RNAs. In other words, reads become oriented such that their poly-A tails are on the right.

### Transcriptome assembly

The transcriptome was assembled using the rnaSPAdes v. 3.15.5 [27] with the following command: *rnaspades*.*py -m 1000 -t 100 -1 Illumina_R1*.*fastq -2 Illumina_R2*.*fastq--nanopore nanopore*.*fastq -o results*.

### The selection of isoforms with longest CDS

ORF prediction was performed using TransDecoder v. 5.5.0 [30] and DIAMOND tool v. 2.1.8.162 [31]. At the first stage, module TransDecoder.LongOrfs with default parameters was used to define the longest ORF. The command used: *TransDecoder*.*LongOrfs -t transcripts*.*fasta*, where *transcripts*.*fasta* is an output of rnaSPAdes. Then, protein sequences from non-redundant NCBI protein database were used to create a database with the command *diamond makedb --in nr --taxonmap prot*.*accession2taxid --taxonnodes nodes*.*dmp -- taxonnames names*.*dmp --db nr* and an output of TransDecoder.LongOrfs (amino acid sequences) were searched against created database with DIAMOND blastp (the command *diamond blastp --query longest_orfs*.*pep --db nr*.*dmnd --max_target_seqs 1 --outfmt 6 -- threads 100 --out blastp*.*outfmt6*). At the last stage ORFs and results of DIAMOND search were used as input for another TransDecoder module, TransDecoder.Predict with parameter “*--single_best_only*” (the command *TransDecoder*.*Predict -t transcripts*.*fasta -- retain_blastp_hits blastp*.*outfmt6 --single_best_only*). After that, for each group (“a gene”) defined by rnaSPAdes the transcript (isoform) with longest protein-coding sequence was retained using a python custom script implemented in the pipeline.

### Transcript clustering

The longest isoforms were clustered using CD-HIT-EST v. 4.8.1 [32] with the following command *cd-hit-est -i cds_longest_iso*.*fasta -o idy_0*.*98_overlap_0*.*6*.*fasta -c 0*.*98-G 0 -aS 0*.*6 -T 32 -p 1 -g 1*, which returns a set of longest isoform from each cluster. A custom script was used to extract corresponding peptides and retain only isoforms with longest CDS in the gff3 annotation file.

### Assembly filtration

The amino acid sequences of the assembly were searched against the non-redundant NCBI protein database (nr.dmnd) using the DIAMOND tool v. 2.1.7.162 [31] with the command *diamond blastp --db nr*.*dmnd --query prots_after_clustered*.*fasta --evalue 1e-5 -- max-target-seqs 8 --threads 100 --out diamond_compairing_res*.*tsv --outfmt 6 qseqid sseqid staxids*. The python custom script implemented in the pipeline was used to identify foreign transcripts. If 50% or more of BLAST hits with taxon IDs (or three or more BLAST hits with taxon IDs) did not belong to the Embryophyta clade, a transcript was considered contaminant and excluded from the assembly.

### Gene Ontology annotation of the transcripts

Gene Ontology annotation of transcriptome assembly was done using PANNZER2 [33]. To obtain GO annotation using Python queries and Beautiful Soup libraries, we uploaded amino acid sequences of transcripts to the PANNZER2 website (http://ekhidna2.biocenter.helsinki.fi/cgi-bin/sans/dump_panz.cgi).

For the emulation of a species from an unexplored taxon all Brassicaceae species were excluded from the protein database downloaded from Phytozome 13 [34] and remaining sequences were used to create DIAMOND blastp database with command *diamond --in phyto_go_db/phyto_prots_for_GO*.*faa --db eggnog_proteins*.*dmnd*. Next, eggNOG-mapper v2 [35] was used to create GO annotation with following commands *download_eggnog_data*.*py -y*, and *emapper*.*py -m diamond --itype proteins -i protein_fasta*.*fasta -o arabidopsis --cpu 30 --dmnd_db eggnog_proteins*.*dmnd*.

### Illumina and ONT read mapping

Illumina paired reads were aligned on the reference *A. thaliana* genome (TAIR10.1) and reads per gene were counted using STAR v. 2.7.10b [36] with parameters *-- sjdbGTFfeatureExon exon --sjdbGTFtagExonParentTranscript transcript_id -- sjdbGTFtagExonParentGene gene_id --quantMode GeneCounts*.

ONT reads were aligned on the reference *A. thaliana* genome (TAIR10.1) using minimap2 v. 2.24 [37] with parameters *-ax splice*.

### The coverage of Arabidopsis genes by assembled transcripts

Assembled transcripts were mapped on the reference *A. thaliana* genome (TAIR10.1) using minimap2 v. 2.24 [37] with parameter *“-x splice”*. Samtools view v. 1.15.1 [38] with parameters *“-Sb”* was used to convert minimap2 output to bam format and bam file was sorted using samtools sort. To match the assembled transcripts with *Arabidopsis* genes bedtools intersect v. 2.30.0 [39] was used with parameters “-wao” and *A. thaliana* genome annotation. The number of transcripts covering each gene was extracted from bedtools intersect output.

### Differential gene expression analysis

Differentially expressed genes were identified with the R package “DESeq2” v. 1.38.3 [40] and thresholds of |logFC| 1 and FDR < 0.05.

### GO enrichment analysis

GO enrichment analysis was performed with the R package “topGO” v. 2.48.0 [41]. P-values of all tests were adjusted via FDR (R “stats” function p.adjust). Terms with FDR < 0.05 and fold enrichment value > 2 were considered statistically significant.

### Pipeline availability

The Trans2express pipeline is available at GitHub via link: https://github.com/albidgy/trans2express. The web-based version of Trans2express is available at http://83.149.250.217/trans2express/.

## Results

### Transcriptome assembly pipeline

The Trans2express pipeline (the overview is depicted at Fig. 1) is based on the rnaSPAdes tool [24] used in a hybrid transcriptome assembly mode. rnaSPAdes demonstrated an excellent performance both in short-read mode benchmark [42] and in comparison of hybrid assemblers [24]. As an output rnaSPAdes generates a set of contigs corresponding to transcripts (mRNA isoforms) grouped by alleged correspondence to a locus.

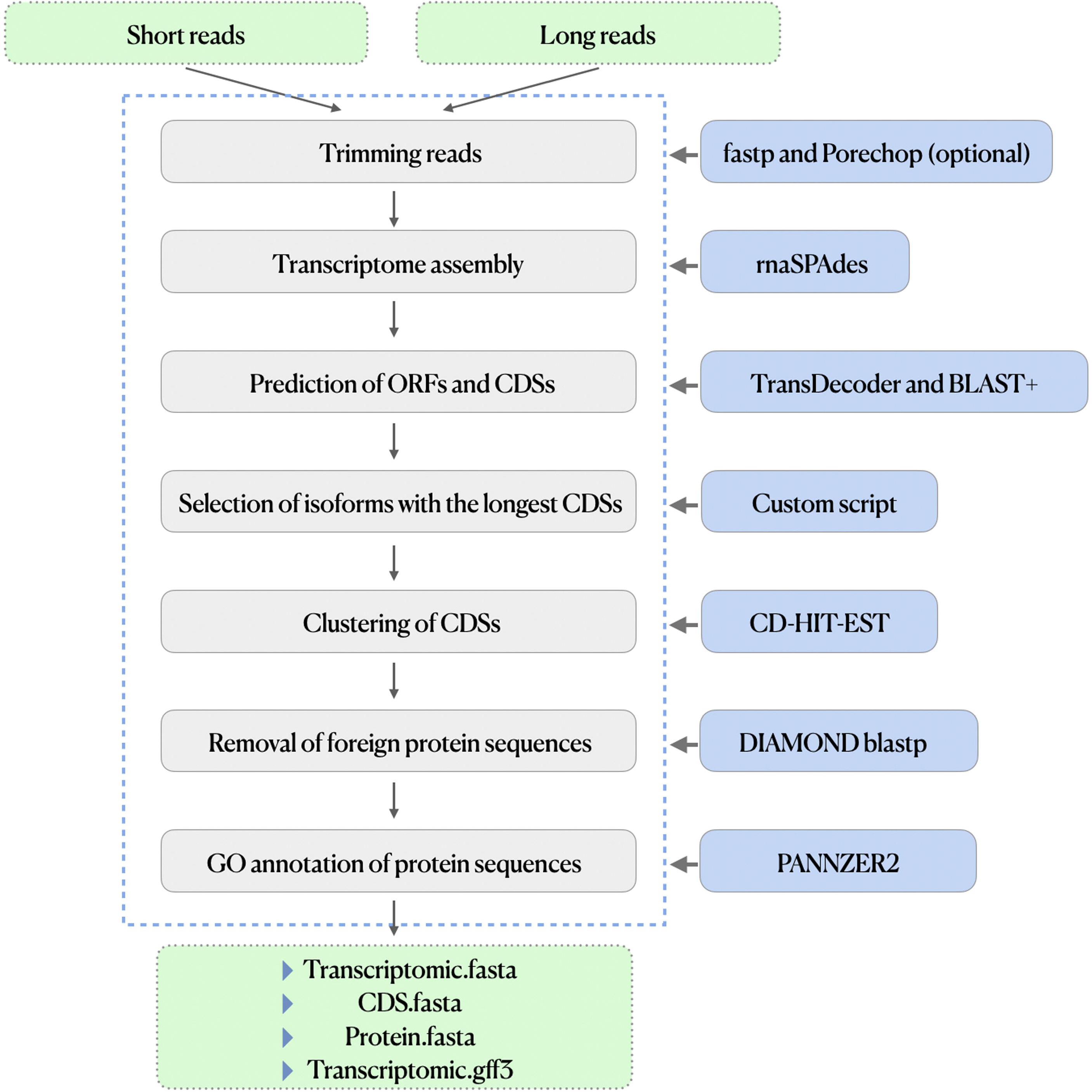

As multiple isoforms in an assembly are known to decrease gene expression analysis performance [16], a transcriptome filtration is a crucial next step. The longest isoform for each group is extracted using TransDecoder [30] and DIAMOND [43]. First, module TransDecoder.LongOrfs with default parameters defines the longest open reading frames (ORFs) of at least 100 amino acids in each transcript. The output amino acid sequences are searched against non-redundant NCBI protein database with DIAMOND and parameter *“-max_target_seqs 1”*. Next, ORFs and results of DIAMOND search are used as an input for another TransDecoder module, TransDecoder.Predict with parameter *“--single_best_only”*. As a result, lists of CDSs and proteins for each transcript are generated. Using a custom script, one transcript with the longest CDS is retained for each rnaSPAdes group.

The next step of assembly filtration includes clustering of transcripts with CD-HIT [32]. CD-HIT performs local alignment and clusters transcripts by sequence similarity. The parameters of CD-HIT are relaxed with respect to similarity and length overlapping in order to increase the isoform number reduction.

At the last step the search of contaminant transcripts is done. If such transcripts are present in the assembly, they will be filtered out. The amino acid sequences corresponding to CDS of the assembly transcripts are searched against the non-redundant NCBI protein database (nr.dmnd, accessed at April 9th, 2023) using blastp of the DIAMOND tool [31].

Taxon IDs of BLAST hits are stored for each transcript; if three or more out of five hits do not belong to Embryophyta clade, a transcript is considered foreign and is removed from the assembly. If less than five hits with taxon IDs are detected, the transcript is also considered as contaminant and removed from the assembly.

After these steps the resulting assembly consists mostly of single transcripts (represented by the longest CDS) per expressed gene of a studied organism.

In order to generate and check biological hypotheses on the process-of-interest it is essential to have functional annotation of the transcripts. Gene Ontology (GO) [44] is the most popular functional classification of genes and plenty of methods for GO annotation inference based on sequence similarity are available so far [45]. In the current pipeline we use command line version of PANNZER2 [33] in order to create a GO annotation of transcriptome assembly.

Trans2express pipeline can be easily download form GitHub (https://github.com/albidgy/trans2express), installed at Linux of MacOS machines and executed from the command line.

### Trans2express online tool

We developed an online version of Trans2express pipeline (http://83.149.250.217/trans2express/). A user should upload short (both single- and paired-end reads are supported) and long reads and provide an e-mail address where notification of the job completion will be sent.

### Assembly of Arabidopsis thaliana transcriptome

To test this pipeline we obtained RNA-seq data of *A. thaliana* Columbia accession (leaves, three replicates). Each replicate was sequenced both on Illumina and Oxford Nanopore platform resulting in 105 M of high-quality short reads and 1.4 M of filtered long reads (Additional file 1: Table S1).

To access the congruence of technologies regarding gene expression Illumina and ONT reads were mapped separately on *Arabidopsis* reference genome (Additional file 1: Table S1). The number of genes that were identified as expressed based on Illumina data was 15 546 (number of reads greater or equal to 16 reads in each replicate, strict threshold [46]), while ONT identified 8 526 expressed genes with threshold one read in each replicate. The majority of genes (97.4%) detected by ONT were also detected in Illumina data and their expression levels were congruent between platforms (Additional file 2: Fig. S1). Genes present in Illumina or ONT data only were mostly expressed at low level and can potentially be absent in sequencing library due to the stochastic effects during library preparation.

ONT and Illumina reads were used for transcriptome assembly via pipeline described above. The primary assembly generated by rnaSPAdes consisted of 55 850 raw contigs (transcripts) with N50 = 2 032 bp which were clustered by rnaSPAdes into 43 054 isoform groups. Raw transcripts were submitted to TransDecoder for ORF annotation, then the longest CDS of each rnaSPAdes-generated isoform group was selected. 19 540 transcripts corresponding to the longest CDSs were combined into a draft transcriptome assembly (N50= 2 017 bp). At the next stage CDSs of the draft transcriptome assembly were clustered by sequence similarity using CD-HIT, which returned the longest CDS of each cluster.

Transcripts of 19 110 CD-HIT-selected CDSs (N50 = 2 036 bp) were subjected to the search of contaminating transcripts. Transcripts with 50% or more non-plant hits in the first five blastp results were filtered out, as well as transcripts containing ambigious nucleotides (N) in their sequences. The final transcriptome assembly included 18 281 transcripts (N50 = 2 051 bp).

### Comparison of reference-based and assembly-based results

In an ideal case, the results of transcriptome assembly of a certain organ/tissue/condition should resemble the results based on a reference genome for the same organ/tissue/condition in terms of the number and content of expressed genes.

At first step we checked the correctness of the transcriptome assembly. In an ideal case the contigs of the assembly should have 1-to-1 correspondence with *Arabidopsis* representative gene models and to align on their sequence on the whole length. Transcripts of both draft (after ORF prediction but before clustering with CD-HIT) and final assemblies were aligned on the *Arabidopsis* reference genome. 96.0% and 99.4% of transcripts for draft and final assemblies, respectively, were successfully mapped on *Arabidopsis* genes. The contigs of the final assembly covered 86.1% of *Arabidopsis* genes expressed in the samples in a single copy (while for the draft assembly before CD-HIT clustering 84.8% of contigs matched a single gene). The maximum number of contigs mapped on one gene (i.e. isoforms) dropped from 11 to 9 (prior to CD-HIT clustering draft assembly comparing with final assembly), indicating the importance of an additional clustering step with CD-HIT.

The identity between translated sequences of *Arabidopsis* CDS and corresponding contigs showed a perfect match for 82.2% of 17 679 gene-contig pairs having identity ≥ 90 (Fig. 2a). The lengths of corresponding proteins arising from transcriptome assembly and reference genome were similar: 87.6% of protein pairs had length ratio close to one (Fig. 2b), indicating on that the contigs represent full-length transcripts.

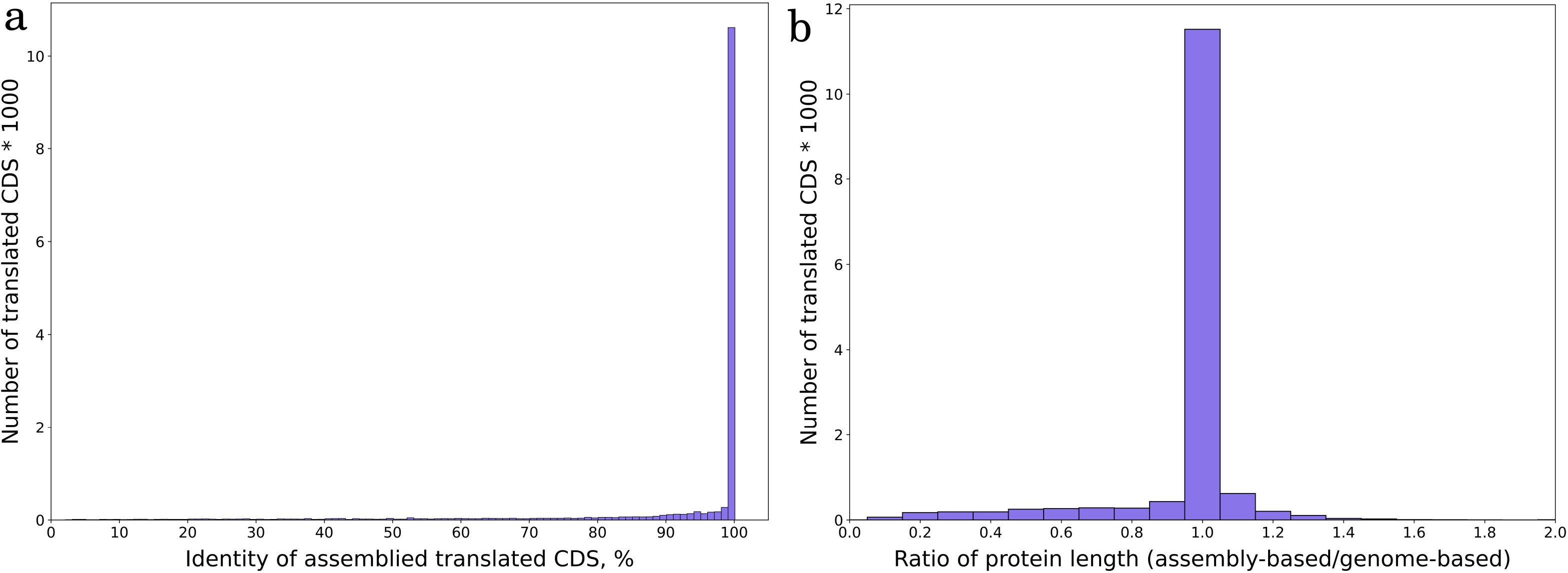

The next step was the functional annotation using Gene Ontology. As Gene Ontology is a hierarchical network, the different GO annotation procedures may result in assignment of GO terms to a gene at different levels of the Ontology tree. To make annotations more comparable, GO annotations for both *A. thaliana* genome and transcriptome assembly were brought to the third level of the Ontology tree counting from the top terms. 17 679 of genes with a single corresponding contig were used in the comparison of GO annotation. 85.6% of transcripts were annotated with at least one GO term. Annotation of 80.7% of contigs at least partially coincided with the annotation of corresponding genes and only 1 196 contigs had annotation that included the same categories as corresponding *Arabidopsis* genes (i.e. the annotation for genes was nested within that of contigs). 14.4% of contigs were not annotated with any GO terms; the majority (92.5%) of the corresponding genes had GO terms assigned and were enriched in growth processes, stress response, cell wall metabolism, ribonucleases, translation, and kinases (Additional file 1: Table S2). GO annotation inferred with exclusion of Brassicales from the protein database (in order to simulate the procedure for a species from a poorly explored clade) returned much poorer results. Only 56.1% of transcripts were annotated with at least one GO term. 45.5% of transcripts had somehow matching with genome ground truth annotation, with 3 174 transcripts having annotation that included the same categories as corresponding *Arabidopsis* genes.

### Gene expression analysis using transcriptome assembly

To explore the applicability of transcriptome assembly for downstream analyses, in particular, for the investigation of gene expression, we performed a model experiment aimed at the investigation of gene expression profiles in response to the change of darkness to light. To do this we collected leaves of *A. thaliana* in two replicates at nine hours and at 21 hours after Zeitgeber (switching the light on, ZT09 and ZT21, respectively). cDNA libraries were sequenced on Illumina, resulting in 37-46M reads (Additional file 1: Table S3). High-quality reads were mapped on *Arabidopsis* genome and on transcriptome assembly. The comparison of expression levels of genes and contigs (based on 17 679 genes with a 1-to-1 correspondence to contigs) revealed high correlation (Pearson r^2^ = 0.77, Fig. 3a).

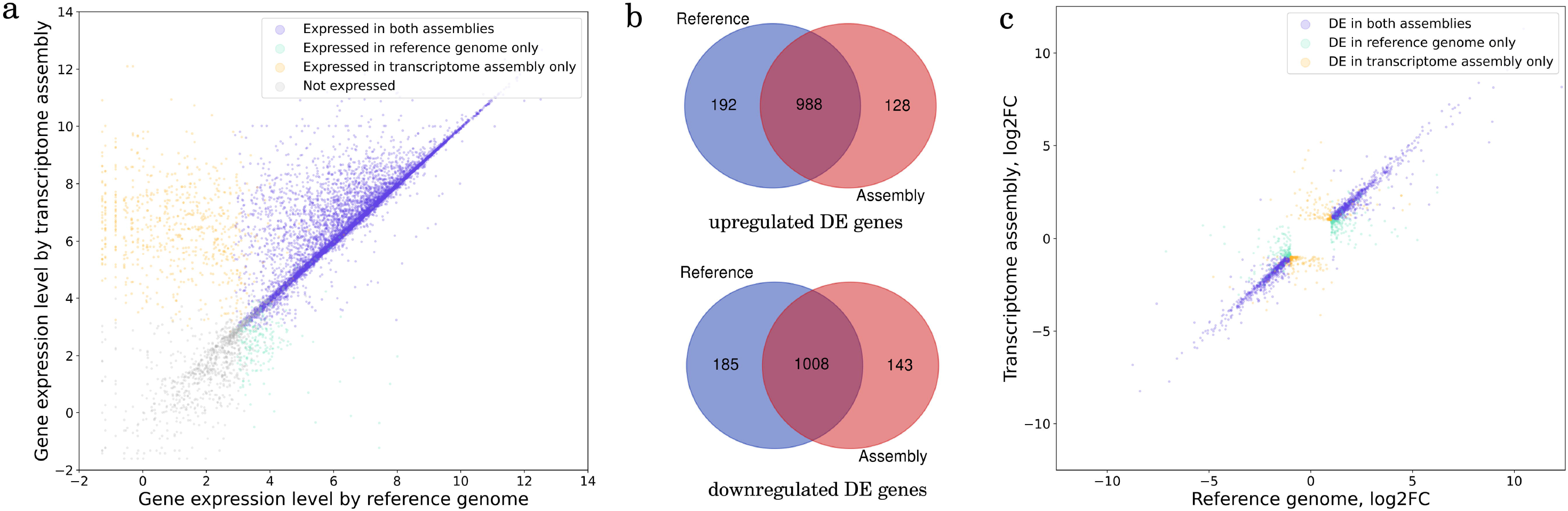

At the next stage, differential expression analysis was conducted using ZT21 (“dark”) as a control sample and ZT09 (“light”) as an “experimental” sample. The number of differentially expressed (DE) genes/contigs was slightly higher for *Arabidopsis* genome than transcriptome assembly: 1 514 and 1 695 genes were up- and downregulated in genome-based results, when transcriptome assembly yielded 1 236 and 1 314 DE genes, respectively. Both up- and down-regulated genes highly overlapped between genome and transcriptome gene lists with 87-88% congruence (Fig. 3b). The majority of DE genes discovered by either reference genome or transcriptome assembly only had fold changes close to the threshold (Fig. 3c) and tend to have lower expression level (Additional file 1: Fig. S2).

Gene Ontology enrichment analysis is a common first step to functional study of expression difference between samples. We conducted GO enrichment analysis for both genome and transcriptome assembly-based DE genes (up- and down-regulated genes were treated separately). Again, genome-based results were wider: 49 terms were found for assembly-based and 210 — for genome-based DE genes; the intersection was 59.2%. Up-regulated assembly-based DE genes had 93 overrepresented terms, and the intersection with 213 genome-based DE genes was 61%. While being less pronounced, GO enrichment of transcriptome assembly-based DE genes still reflects the main processes associated with circadian cycle and difference between light and dark plant metabolism (Additional file 1: Table S4).

## Discussion

Non-model species, especially plants, often bear large and complex genomes demanding a great deal of resources for sequencing and assembly. Fortunately, a wide set of scientific problems may be addressed with sequences of RNAs, i.e. transcriptome. Among a large variety of *de novo* assembly approaches the hybrid assembly utilizing accurate short reads and transcript-spanning but less accurate long reads is a state-of-the-art approach resulting in high quality transcripts [24].

Many eukaryotic genes are prone to alternative splicing that generates several isoforms per locus. Theoretically *de novo* transcriptome assembly has potential to recognize all expressed isoforms of a gene, and some approaches (e.g. RATTLE [47] or isONform [48] are aimed to reconstruct as many isoforms as possible. However, for many types of downstream analyses the uncollapsed isoforms are disadvantageous (the same is true for allelic variants of transcripts). In case of the use of assembly for evolution-centered studies (phylogenetic analysis, investigation of gene families) isoforms or alleles can be incorrectly recognized as paralogs and thus lead to biases and incorrect biological conclusions. In case of gene expression analysis they will adversely affect the estimation of gene expression levels as many RNA-seq reads will be non-uniquely mapped resulting in a loss of data [16]. Thus, an extensive clustering and filtration of *de novo* transcriptome assembly are crucial steps. There are several pipelines that address this problem, for example based on the minimization of the fraction of duplicated genes in BUSCO gene sets [49]. This approach however might not be applicable for polyploid species (that are especially frequent in plant lineage) because it will collapse paralogs.

Here we present Trans2express – a pipeline for *de novo* transcriptome assembly optimized for one of the most frequent downstream applications – gene expression analysis. We tested our approach on *A. thaliana* leaf transcriptome sequenced on Illumina and ONT platforms. Almost all transcripts of the assembly were mapped on the *Arabidopsis* genome and the quality of transcripts was high in terms of protein identity and length ratio. 86.1% of expressed genes were covered by a single transcript, in contrast to what is typical for transcriptome assemblies.

The congruence of the results of differential expression analysis between reference-based and assembly-based variants indicates on the high potential of the pipeline for gene expression analysis in non-model species. It is important to note that the discrepancies that are still present between reference-based and assembly-based analyses are confined to the genes with lower expression values and with fold change close to the threshold (logFC = 1). From the practical point of view this means that when dealing with the results of gene expression analysis based on *de novo* assembly it might be beneficial to set stricter thresholds on FC. This will recover less DE genes but with higher reliability.

The integral part of transcriptome characterization and the differential gene expression analysis is Gene Ontology (GO) annotation and further search of the enrichment of specific GO terms in DE genes. Despite that, quite few studies focus on (or at least discuss) the issues of GO annotation in non-model species. We show here that this step produces very dissimilar results depending of the datasets taken as a basis for homology search and does not work perfectly even for a model species. Indeed, surprisingly, even if the information on *Arabidopsis* is not excluded, the GO annotation of *Arabidopsis* contigs yields quite poor results – only 80.7% had annotations overlapping with the reference ones and ∼ 14% remained unannotated. Under the simulation of the case of non-model species which was done by exclusion of information of *Arabidopsis* and other Brassicaceae only 56.1% annotations overlapped with the reference ones. This highlights the challenge of the GO-annotation in non-model species and calls for the development of more model species representing different plant lineages. Any analysis dealing with GO in non-model species should be treated with caution, especially if it involves cross-species comparisons. Despite these considerations, in case of strong and pronounced effect the results will be biologically relevant, as shows the model experiment done in this study.

## Conclusion

Despite the existence of many pipelines for transcriptome assembly and filtering few of them were tested in downstream applications. Here we present Trans2express – a procedure that allows to get a de novo assembly out of combination of short and long reads optimized for the generation of a single isoform per gene. This is important for many applications, including the search of differential gene expression. Using experiments on model plant *Arabidopsis thaliana* we showed that our pipeline allows the assembly of more than 85% of expressed genes in a single isoform. The differential expression analysis showed high concordance between reference-based and *de novo* transcriptome assembly-based results. We expect that the method described here as well as online tool will promote studies on plant species with large genomes, where the transcriptome assembly is a feasible alternative to the genome assembly.

## Declarations

### Ethics approval and consent to participate

Not applicable.

### Consent for publication

Not applicable.

### Availability of data and materials

The *Arabidopsis de novo* transcriptome assembly is available at 10.6084/m9.figshare.24981996. The Illumina and ONT reads used for *de novo* transcriptome assembly are available at SRA NCBI database (https://www.ncbi.nlm.nih.gov/sra/) under BioProject accession PRJNA1063591. The Illumina reads used for gene expression analysis are available at SRA NCBI database (https://www.ncbi.nlm.nih.gov/sra/) under BioProject accession PRJNA1063592.

Trans2express pipeline is available at GitHub via link https://github.com/albidgy/trans2express.

### Authors’ contributions

AVK and AMK developed pipeline and analyzed the data, ASK developed initial version of pipeline, AAP and MDL conceived and coordinated the study and obtained experimental data, MIS tested the pipeline and developed online version, AVK and MDL wrote the article.

## Acknowledgements

Not applicable.

