## Additional file 2 for "Trans2express – de novo transcriptome assembly pipeline optimized for gene expression analysis"

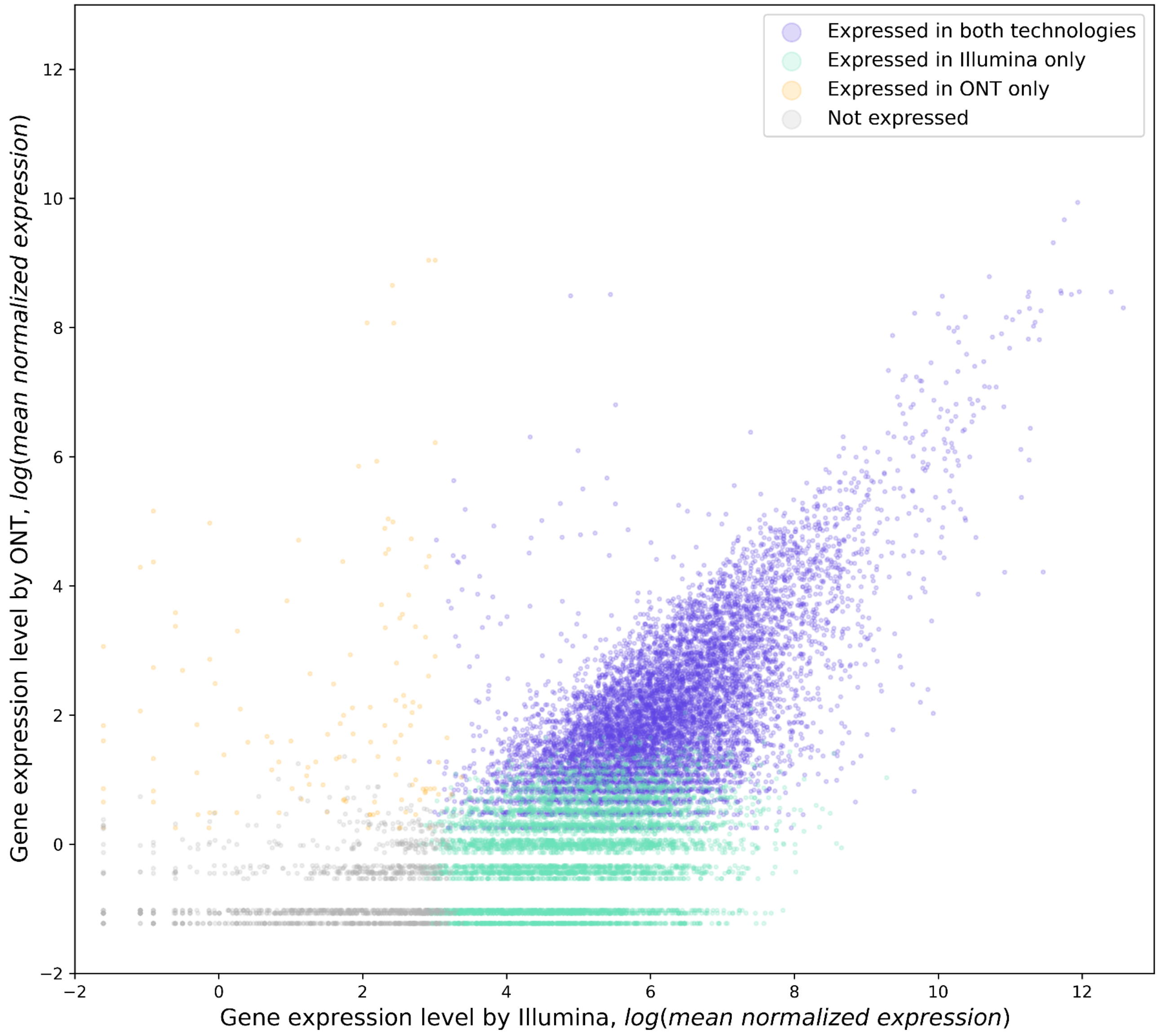


Fig. S1. Congruence of gene expression level inferred by ONT and Illumina sequencing. Genes, identified as expressed by both technologies, by ONT or by Illumina only are marked.

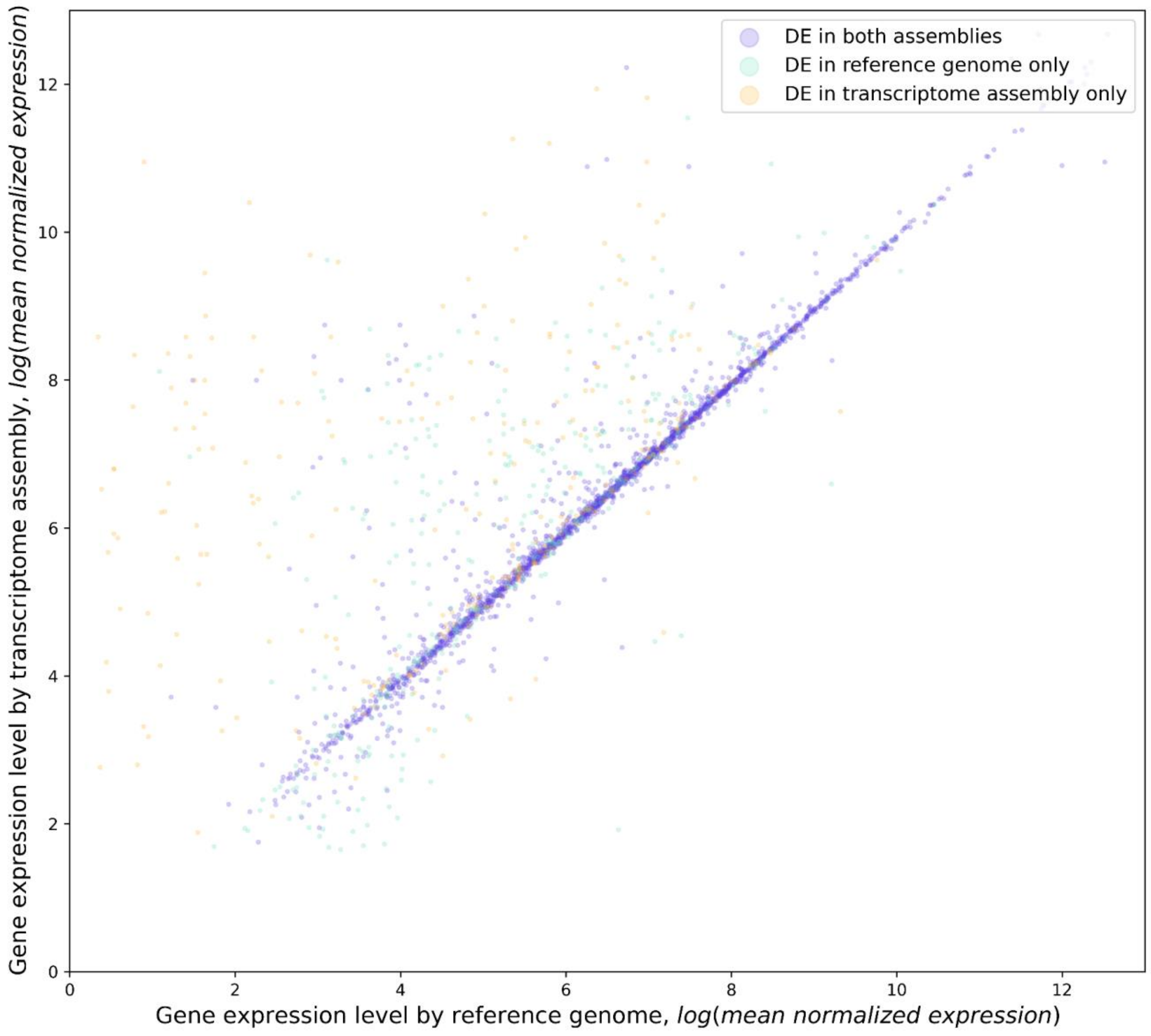


Fig. S2. Expression level of differentially expressed genes identified using genome reference and transcriptome assembly. Genes, identified as expressed by both approaches, by genome or by transcriptome assembly only are marked.
